# Disentangling non-random structure from random placement when estimating β-diversity through space or time

**DOI:** 10.1101/2023.09.19.558467

**Authors:** Daniel J. McGlinn, Shane A. Blowes, Maria Dornelas, Thore Engel, Inês S. Martins, Hideyasu Shimadzu, Nicholas J. Gotelli, Anne Magurran, Brian McGill, Jonathan M. Chase

**Author notes:** Corresponding & contact author.

## Abstract

There is considerable interest in understanding patterns of β-diversity that measure the amount of change in species composition through space or time. Most hypotheses for β-diversity evoke nonrandom processes that generate spatial and temporal within species aggregation; however, β-diversity can also be driven by random sampling processes. Here, we describe a framework based on rarefaction curves that quantifies the non-random contribution of species compositional differences across samples to β-diversity. We isolate the effect of within-species spatial or temporal aggregation on beta-diversity using a coverage standardized metric of β-diversity (β_C_). We demonstrate the utility of our framework using simulations and an empirical case study examining variation in avian species composition through space and time in engineered versus natural riparian areas. The primary strengths of our approach are that it provides an intuitive visual null model for expected patterns of biodiversity under random sampling that allows integrating analyses across α-, γ-, and β-scales. Importantly, the method can accommodate comparisons between communities with different species pool sizes, and can be used to examine species turnover both within and between meta-communities.

**Open Research statement:** all code and data used in this manuscript are available at the following link: https://github.com/MoBiodiv/beta_concept

## Introduction

Ecologists are frequently interested in how the composition of species in a community changes across space or time (Scheiner et al. 2011, Magurran et al. 2019, Daskalova et al. 2020).The degree of change in species composition in assemblages across space or time is often referred to as β-diversity: localities or time periods with fewer species in common have higher β-diversity. Most conceptual explanations of β-diversity evoke processes that generate non-random spatial or temporal patterns of species aggregation (Leibold and Chase 2018). Aggregation here refers to clustering whereby individuals occur near other individuals of the same species in time and/or space. For instance, two of the most commonly discussed mechanisms underlying patterns of β-diversity are environmental filtering and dispersal limitation (Legendre et al. 2005, Vellend 2016, Leibold and Chase 2018). Both of these mechanisms increase aggregation of species distributions via conspecific clustering in space or time increasing β-diversity.

Although most attention has been focused on the non-random mechanisms underpinning β-diversity, it can also reflect random sampling effects of individuals and species taken from multiple points in space or time. Imagine we collect a sample of 40 individuals within a region or time period that supports up to 50 different species. Even in the improbable case that the numbers of individuals of each species are exactly the same (completely even), at least 10 species will be excluded from our sample because of the limited number of individuals. If we then compare that sample to another from a different location in space or time, a different set of 10 (or more) species will be excluded simply due to random sampling effects: the species composition of the two samples will differ entirely due to incomplete sampling. This phenomenon has been variously termed a “sampling effect” (e.g., Adler et al. 2005), a “rarefaction effect” (e.g., Palmer et al. 2008), and the “random placement model” (e.g., Coleman et al. 1982). The core idea is that the number of species observed in a sample is constrained by the number of individuals in that sample. Returning to our thought experiment, if the species have a more realistic abundance distribution, with many individuals of a few common species and many species with few individuals (i.e., rare species), these sampling effects on β diversity can be strong (Kraft et al. 2011, Chase et al. 2018, McGlinn et al. 2019, Engel et al. 2021). This example emphasizes that spatial or temporal β diversity is potentially underlain by two factors: a) the non-random turnover of species, due to ecological mechanisms such as environmental filtering or dispersal limitation; and, b) the random turnover of species due to incomplete sampling, especially of rare species (i.e., sampling effects).

Most metrics of β-diversity conflate variation from both random sampling effects and spatially non-random mechanisms (Stegen et al. 2013, Chase et al. 2018, McGlinn et al. 2019, Engel et al. 2021, Chao et al. 2023). This means that the same observed change in β-diversity may be due to different underlying mechanisms, sometimes referred to as a “many-to-one problem”, which are common in ecological studies (Frank 2014, Scholes 2017). Specifically, random turnover can occur where there are changes/differences in other non-spatial components of diversity, such as the species abundance distribution and size of the regional species pool and the total number of individuals. To illustrate this, consider the three hypothetical scenarios in Figure 1 using Whittaker’s (1960) β-diversity (*β_S_* = *γ* / *α̅*, where *γ* is the regional, and *α̅* is the average of local diversity). In each scenario (Fig1. a-c), a shift in a different component of community structure results in a doubling of Whitaker’s β-diversity (from 1 to 2). In the first two cases (Fig1.a, b), β-diversity increases due simply to random placement of individuals resulting either from a shift in the species-abundance distribution (SAD), for example, a decrease in evenness (Fig. 1a) or, from a decrease in the total number of individuals (*N*) (Fig. 1b). In the third case, the same magnitude of shift in *β_S_* is due to an increase in conspecific aggregation (Fig. 1c). This “many-to-one” effect is particularly problematic when trying to link changes in β-diversity to hypotheses that evoke changes to conspecific clustering due to environmental filtering or dispersal limitation. To link these mechanisms to β-diversity, it would make sense to focus on patterns of β-diversity that reflect only changes in conspecific aggregation rather than changes in *N* or the SAD (which we refer to as sampling effects). One consequence of β-diversity metrics confounding both random and non-random variation is that most β-diversity metrics can increase as aggregation decreases if *N* is decreasing or the SAD is becoming less even for example.

**Figure 1.**
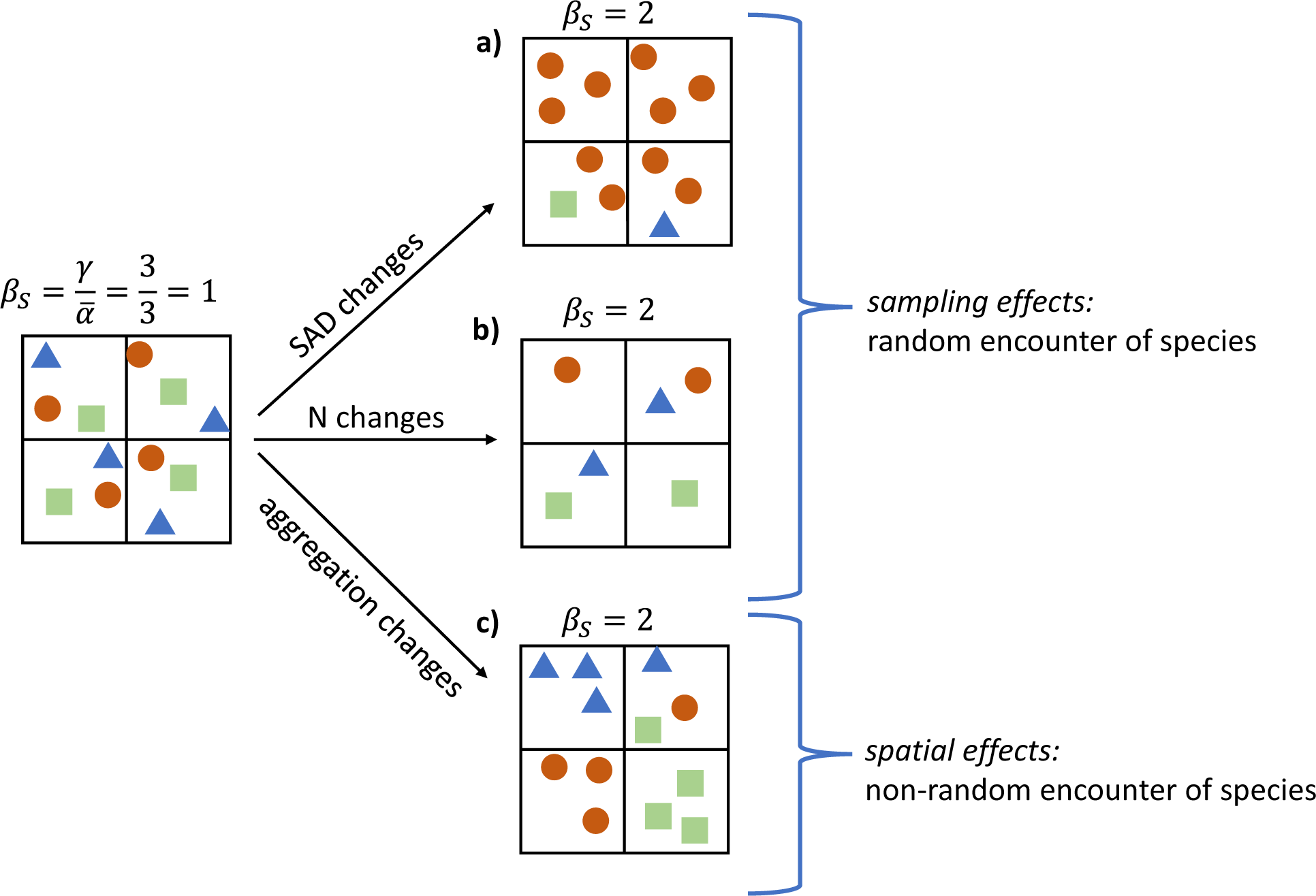
Cartoon communities that illustrate how random sampling effects and non-random spatial effects can result in identical values of Whittaker’s β-diversity (*β*_S_), where *α̅* is average sample richness across plots (small boxes) and *γ* is total species richness in a site (large boxes). The different symbols represent individuals of different species. Panels (a) and (b) illustrate changes in community structure that are consistent with a random sampling model in which spatial β increases either because the regional species-abundance distribution (SAD) is less even (a) or because there are many fewer individuals (b). Panel (c), illustrates how a spatially non-random process such as environmental filtering results in conspecific aggregation producing an identical value of *β*_S_.

It is important to emphasize that this is a problem that potentially influences all β-diversity metrics. Any metric of β-diversity that does not explicitly consider the process of sampling is sensitive to sampling effects. So regardless of whether turnover is calculated using presence-absence vs abundance data, is examined in space or time, or using pairwise vs multisite metrics, if the goal of the analysis is to link patterns of compositional change to mechanisms that generate non-random conspecific occurrence patterns, then sampling effects should be controlled for in the measurement of β-diversity. Other authors have recognized this and proposed a randomization algorithm to try to control for sampling effects on β-diversity (Kraft et al. 2011, Chase et al. 2011, Myers et al. 2013, 2015). Yet, continued debate as to exactly how to develop those randomizations, and just what the deviations mean (Kraft et al. 2012, Qian et al. 2012, 2013, Xu et al. 2015, Tucker et al. 2016) indicates that a more general solution is necessary.

In this paper, we describe a framework for quantifying the non-random contribution of species compositional differences across samples to β-diversity. This framework can be applied to any question related to measuring compositional variation (i.e., β-diversity) across samples, whether it be within a given (relatively homogeneous) metacommunity, across an environmental gradient, or through time. The approach allows us to differentiate the contribution of non-random species compositional shifts from the effects of sampling properties due to random placement to changes in β-diversity. As a result, we can quantify and compare compositional shifts among samples through space or time, and potentially relate these to other features of the system (e.g., changing spatial or environmental conditions). Here, our primary purpose is not to review and/or unify all metrics and measures of β-diversity, nor to advocate for a single superior metric, both of which have been attempted (Tuomisto 2010, Chao et al. 2012, 2023). Rather, we promote a framework for measuring the relative influence of sampling and non-random associations that underlie β-diversity among samples, regardless of whether it is measured within or across landscapes, through time, or any combination thereof. Furthermore, rather than using different concepts and tools, we show how a single conceptual framework can identify the key components underlying variation in species composition.

First, we describe a simple framework that uses rarefaction curves to decompose β-diversity into components due to sampling effects, and those that are due to non-random aggregations of species. Second, we show the framework can be applied to multiple, related questions about how species composition varies across samples.

### A unified framework for dissecting the non-random contribution of species compositional variation to β-diversity in space and time

The components of our framework are not new. The framework is based on a long history of rarefaction and accumulation curves that depict how species numbers increase with increasing sampling effort (Preston 1960, Sanders 1968). For example, Kobayashi (1982, 1983) showed how spatial aggregation could be quantified from rarefaction curves by comparing subsets of spatially explicit samples to the entire range of spatially randomized samples. Likewise, Gotelli and Colwell (2001) showed how comparing accumulation or ‘collectors’ curves that retain spatial information about the distributions of individuals to individual-based rarefaction curves could provide an indicator of the degree to which aggregation influenced spatial patterns of species accumulations (see also Crist and Veech 2006, Chiarucci et al. 2009, Cayuela et al. 2015, Chase et al. 2018, McGlinn et al. 2019). Finally, Olszewski (2004) explicitly discussed how the comparisons between spatially explicit and randomized rarefaction curves could be used as an index of β-diversity (see also Crist and Veech 2006, Dauby and Hardy 2012). These perspectives have been more recently formalized using individual-based rarefaction curves (and related diversity curves) to disentangle non-random structure from random placement underlying β-diversity within a given set of environmental conditions (i.e., a metacommunity) (Chase et al. 2018, McGlinn et al. 2019, 2021, Engel et al. 2021). Here, we generalize this approach and apply it to questions examining β-diversity among different kinds of samples, such as sites across a strong environmental gradient, or when quantifying temporal β-diversity.

Our framework is designed for one of the most common data types available to community ecologists - a sample-by-species matrix. Each sample contains a vector of abundances of all species sampled from a given assemblage and comes from a given local site. Samples can be collected across multiple sites (a site-by-species matrix) or across multiple time periods (a time-by-species matrix), or a combination of the two. For simplicity, we illustrate the different steps of the approach with samples taken from two spatial locations or time points in Figure 2, but it can be generalized to any number of samples. We assume that the communities being compared are sampled in such a way that they have the same sample effort, i.e., the spatial and temporal grain, extent, and sample arrangement are equal across communities (or can be standardized to such). Here, we define a single sample as the α-scale, and the sum of samples as the γ-scale; however, other accumulation schemes are also possible, so long as the α-scale is a subset of the γ-scale.

**Figure 2:**
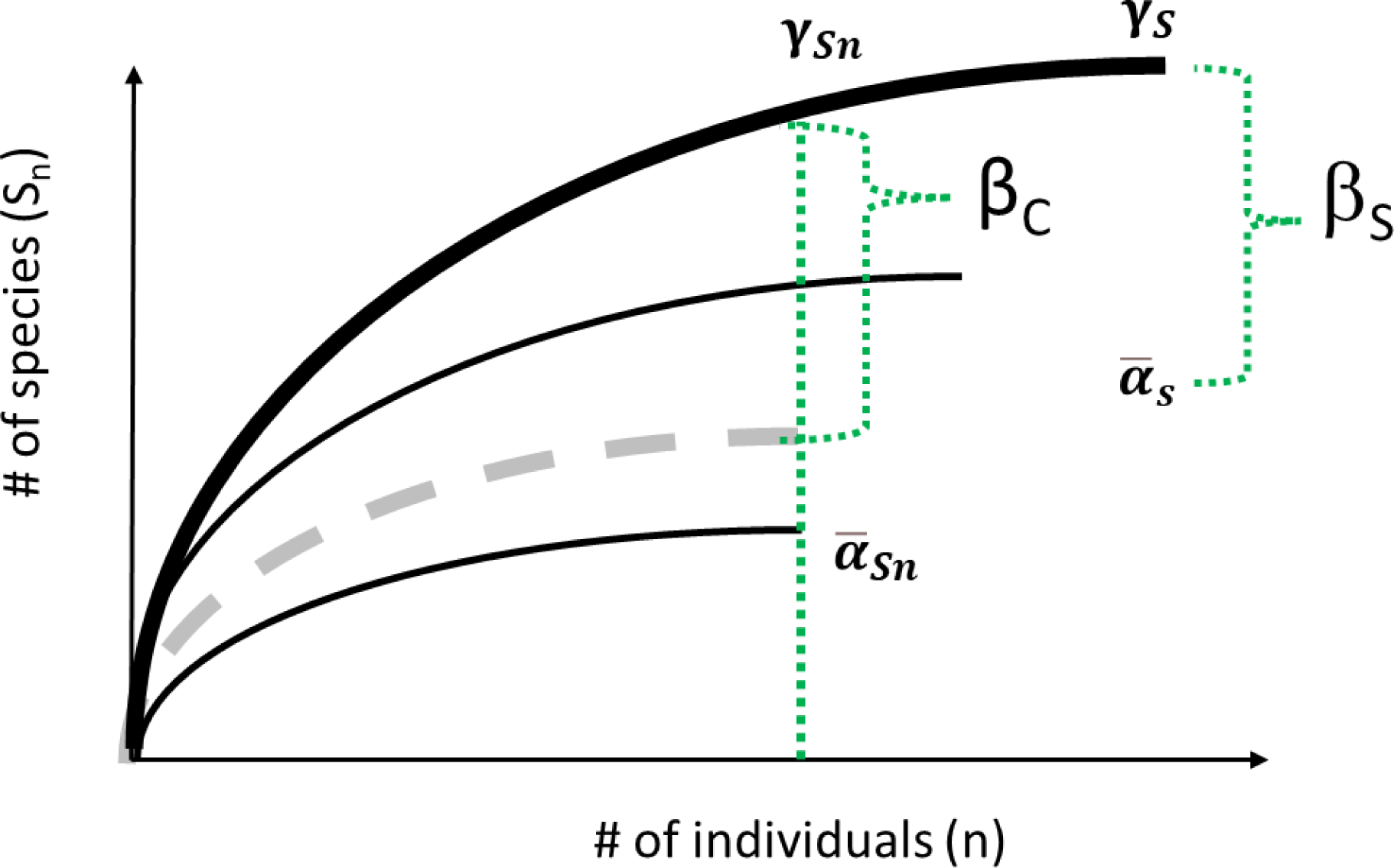
Individual based rarefaction curves at the *α*- and *γ*-scales. Thin solid black lines depict two α-scale curves, with their average shown by the thick dashed gray line; solid thick black line shows the *γ*-scale curve, which is calculated by pooling the two *α*-scale samples. Whittaker’s *β*_S_ is calculated as the ratio of *γ_S_* and *α̅*_***s***_. In contrast, coverage-based *β*_C_ controls for the number of individuals by using the ratio of *γ_Sn_* and *α̅*_***sn***_, where *n* provides a target degree of coverage that adjusts for variation in the regional SAD (species pool) when comparing between meta-communities (eqn 5 from Engel et al. 2021).

#### Step 1: Create a rarefaction curve for the sum of all samples: γ-scale

If we pool the two (or more) samples, we can calculate the γ-scale rarefaction curve (solid, thick black line in Figure 2). This curve shows the number of species for a random sample of individuals from the whole metacommunity or time series; for any sample of *n* individuals, *S_n_* is the expected number of species in that sample. This type of rarefaction curve is sometimes referred to as an individual-based rarefaction curve or random sampling model. The curve and its variance has been derived analytically for sampling without replacement (Hurlbert 1971, see Coleman et al. 1982 for formulation for sampling with replacement). Here, because we calculate rarefaction using all samples, the γ-scale curve represents a ‘null expectation’ of the number of species for *n* individuals, when all individuals of all species occur randomly across the samples (in space or time).

#### Step 2: Create a rarefaction curve for each individual sample and average them: α-scale

Next, we calculate the rarefaction curves for the individual samples (Fig 2, thin solid black lines) and average them to obtain the α-scale rarefaction curve up to the number of individuals (*n*) that provides the target level of coverage (Fig. 2, thick dashed gray line; Engel et al. 2021). Here coverage refers to how close the γ curve has come to a hypothetical asymptote (i.e., it is an estimate of sample completeness, Chao and Jost 2012).

#### Step 3: Compare the α and γ-scale curves to estimate the β-scale patterns

The classical Whittaker’s β_S_ metric is calculated as γ / *α̅* where *α̅* is average sample richness (Whittaker 1960). Within our framework, these values are represented by the ends of the rarefaction curves (i.e., the average number of species per site [*α̅*], and all species observed across all sites in a region or time points [*γ*], Fig. 2).

To estimate non-random spatial structure, we must compare the (average) α-scale curve (dashed line) to the γ-scale curve, after standardizing for sampling effects. We control for the numbers of individuals sampled (i.e., sampling effects) by comparing the γ- and α-scale rarefaction curves at the same value of number of individuals (*n*).

To illustrate the behavior of these metrics, we simulated four scenarios similar to those shown on Figure 1, and calculated β_S_ and β_C_. All scenarios have 50 species in the regional pool, but they vary from the starting community in either their evenness, total number of individuals, or conspecific aggregation. When individuals of all species are distributed randomly and only the evenness of the SAD decreases (Fig. 3A: high evenness 3B: low evenness) or the total number of individuals decreases (Figure 3A: N = 1000; 3C: N = 400), we see that the average *α*-scale curve (dashed gray line) falls directly on top of the γ-scale curve (solid black line, Figure 3 E-G), and low evenness and fewer individuals are associated with an increases in β_S_, but β_C_ is equal to one in both cases (insets on Figure 3F and G). However, when we add non-random structure via species aggregation (Figure 3D), the α- and γ-scale IBR curves diverge (Figure 3H), and both metrics are greater than one (compare inset Figure 3H to 3E).

**Figure 3:**
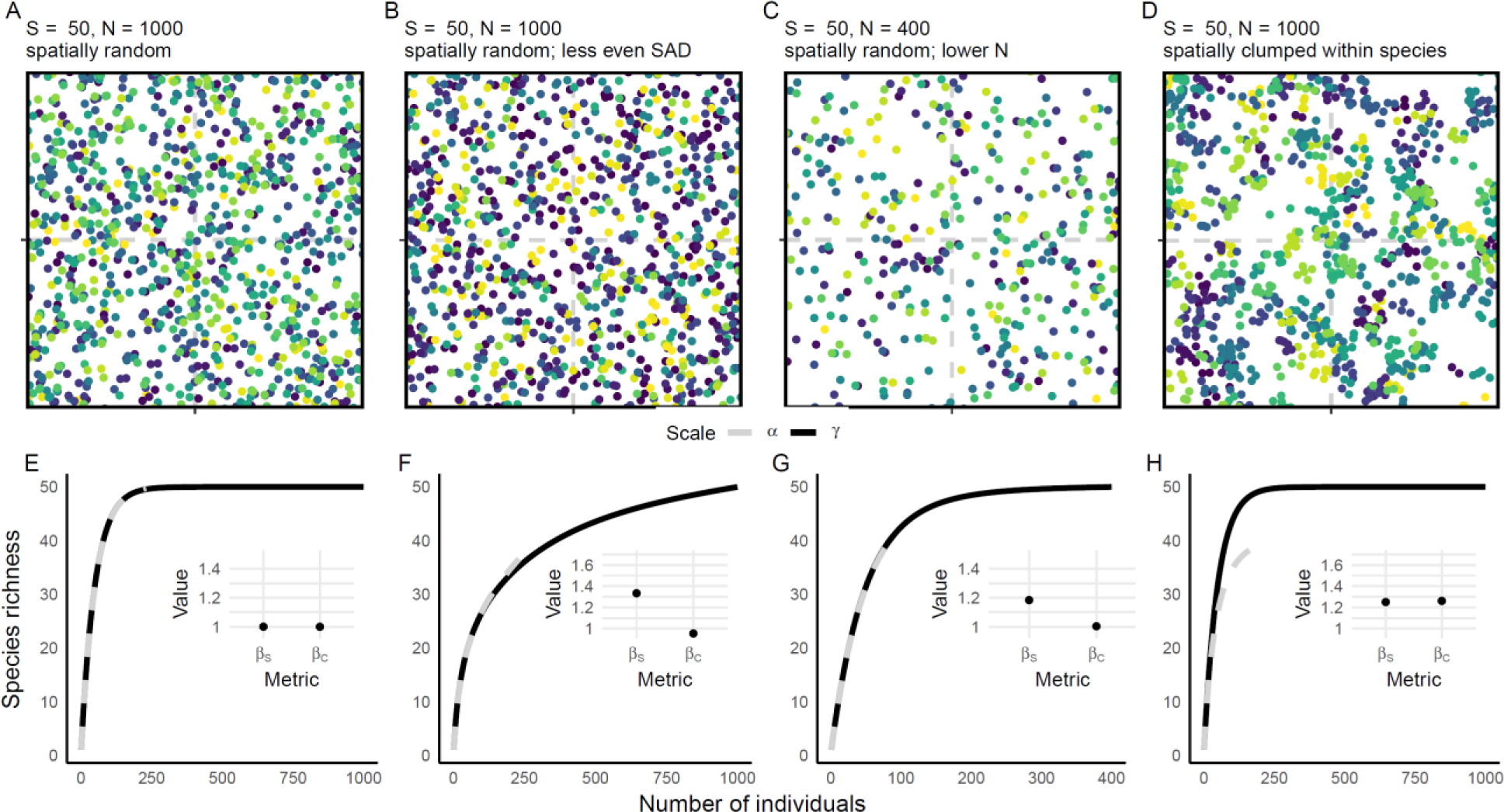
Quantitative illustration that *β_S_* responds to changes to changes in evenness and the total number of individuals, whereas *β_C_* only responds to changes in within-species aggregation. Simulated communities are shown in panels (A-D) in which different colored dots represent individuals of different species. The landscape (*γ* scale) is divided into four quadrats (*α* scale, dashed gray lines). Panels E-H show the corresponding rarefaction curves associated with each artificial community for the *γ*-(solid black line) and average α-scales (dashed gray line). Inset on these panels is the value of each *β* diversity metric described. Note that *β* = 1 means species composition does not vary among samples.

If species are randomly distributed among sites (or time points; Fig. 3A-C), then species will likely be sampled at all sites, and the α- and γ-scale curves will fall on top of each other (Fig. 3E-G). However, if species display conspecific aggregation (i.e., individuals within a species are clumped, Fig. 3D) such that they are non-randomly distributed in space or time, then the α-scale curve will fall below the γ-scale curve (Fig. 3H), because new species will be encountered across different sites or time points due to the within species aggregation, pulling the γ-scale curve up relative to the α-scale curve. The resulting ratio of γ*_Sn_* and *α̅*_*sn*_, which we call coverage-based β diversity (*β_C_,* Engel et al. 2021), is indicative of the degree to which species show (non-random) intraspecific aggregation among sites or time periods.

Thus, β_C_ reflects the degree of non-randomness in the spatial or temporal distribution of species within the domain of (0, ∞). The example in Figure 3D shows a case where there is a non-random distribution of species composition among samples, and β_C_>1. However, it is also plausible that the α- and γ-scale curves could completely overlap (see e.g., Fig. 3E-G), in which case we would conclude that even though there is β-diversity (i.e., β_S_>1), this is simply due to random placement effects (β_C_=1). Finally, species can also show conspecific segregation (i.e., individuals within a species are overdispersed more than random), where the α-scale curve falls above the γ-scale curve, and *β_C_* < 1 (not shown).

One additional benefit of βc (that is not illustrated in Figure 3 but described in detail in Engel et al. 2021) is that this metric is unbiased when comparing β-diversity across meta-communities that differ in the size of their species pools (e.g., in temperate vs. tropical environments, or across strong environmental gradients). This is accomplished by computing β_C_ within each meta-community at the same level of sample coverage or completeness (Chao and Jost 2012). In effect, ensuring that γ-scale sample coverage is the same for all metacommunities means that the value of *n* (the number of individuals for which γ*_Sn_* and *α̅*_*sn*_ are calculated) varies among metacommunities.

To summarize, traditional measures of variation in species composition across area or time (β_S_) are shaped by both random and non-random sampling processes, and we can isolate the non-random structure in space or time in determining that scaling by calculating β_C_ (Table 1). Furthermore, we can evaluate these β-diversity measures for a wide variety of questions concerning species compositional shifts in space and time. We provide R code to calculate classical β_S_ and β_C_ (as well as several other β metrics which we do not show here for simplicity) in mobr∷calc_beta_div (McGlinn et al. 2022).

**Table 1.** Multiplicative *β* diversity metrics and the effects that they capture. Species abundance distribution (SAD) effects are due to changes in species evenness and/or the size of the species pool, N effects refer to changes in richness due to variation in the number of individuals sampled, while aggregation effects refer to changes in richness due to variation in how individuals are spatially or temporally distributed (clumped, random, or overdispersed).

| Metric | Number of individuals sampled ($n$) | Effect controlled for | Effects captured |
| --- | --- | --- | --- |
| $\beta_s$<br>(Whittaker's) | Regional $n$ vs average local $n$ | none | SAD, N, and aggregation |
| $\beta_c$<br>(coverage) | Regional $n$ equals local $n$ and corresponds to a target level of coverage. Across meta-communities, coverage is fixed but $n$ may vary. | N and SAD (evenness and size of pool) | aggregation |

### One approach, many questions: Some example applications

There are several benefits to our approach. Rarefaction curves provide an intuitive visualization of *α*- and *γ*-diversity patterns, the shape of the SAD, and the degree of variation in species composition that exists between samples. Moreover, the same family of measures can be used to estimate *β*-diversity, and to differentiate between random placement and non-random structure leading to biodiversity scaling for multiple related questions. We illustrate some of this potential using a case study. We examined compositional variation in bird diversity between natural and engineered riparian habitats using a subset of data from the Central Arizona-Phoenix Long-Term Ecological Research site (Warren et al. 2022). We focus on riparian habitats where water permanence was perennial, and contrast sites in engineered settings (including a landscaped riparian preserve, a constructed wetland, and a water retention area along the Salt River, each surrounded by urban or agricultural areas) with those in more natural environments (located along perennial river reaches and surrounded by desert). Point count surveys with a 40-m fixed radius were conducted by trained observers that recorded all birds seen and heard; we analyzed samples collected in spring between 2001 and 2016. Before calculating our metrics, we ensured that sample effort was consistent across all sites and years; this meant three sites were retained from each habitat (engineered and natural), and data from 2003 and 2009 were discarded due to missing samples.

Using these effort-standardized data, we address four questions examining how random and non-random components contribute to patterns of β-diversity through space and time: (1) does the total spatiotemporal variation in community composition differ between engineered and natural habitats? (2) How does spatial variation in community composition change through time in each of the two habitats? (3) Does the temporal variation in community composition differ between engineered and natural habitats? (4) Are there compositional differences between (rather than within) engineered and natural habitats, and do any differences change through time?

#### Q1) does the total spatiotemporal variation in community composition differ between engineered and natural habitats?

We used all site-year combinations within each habitat to examine total spatiotemporal variation in community composition. γ-scale rarefaction curves combine all the samples across space and time within habitats, and show that the engineered habitat had more individuals, but fewer species than the natural habitat (Figure 4i). To examine spatiotemporal variation, we defined the α-scale as a single site-year combination within a habitat (Figure 4i inset shows α- and γ-scale curves). The greater number of species in the natural habitat compared to the engineered habitat resulted in higher β*_S_* in natural habitats. However, this pattern was reversed for β_C_ when the influence of sampling effects were removed from the calculations (Figure 4ii), meaning that aggregation in time and space was similar in the engineered and natural habitats.

**Figure 4.**
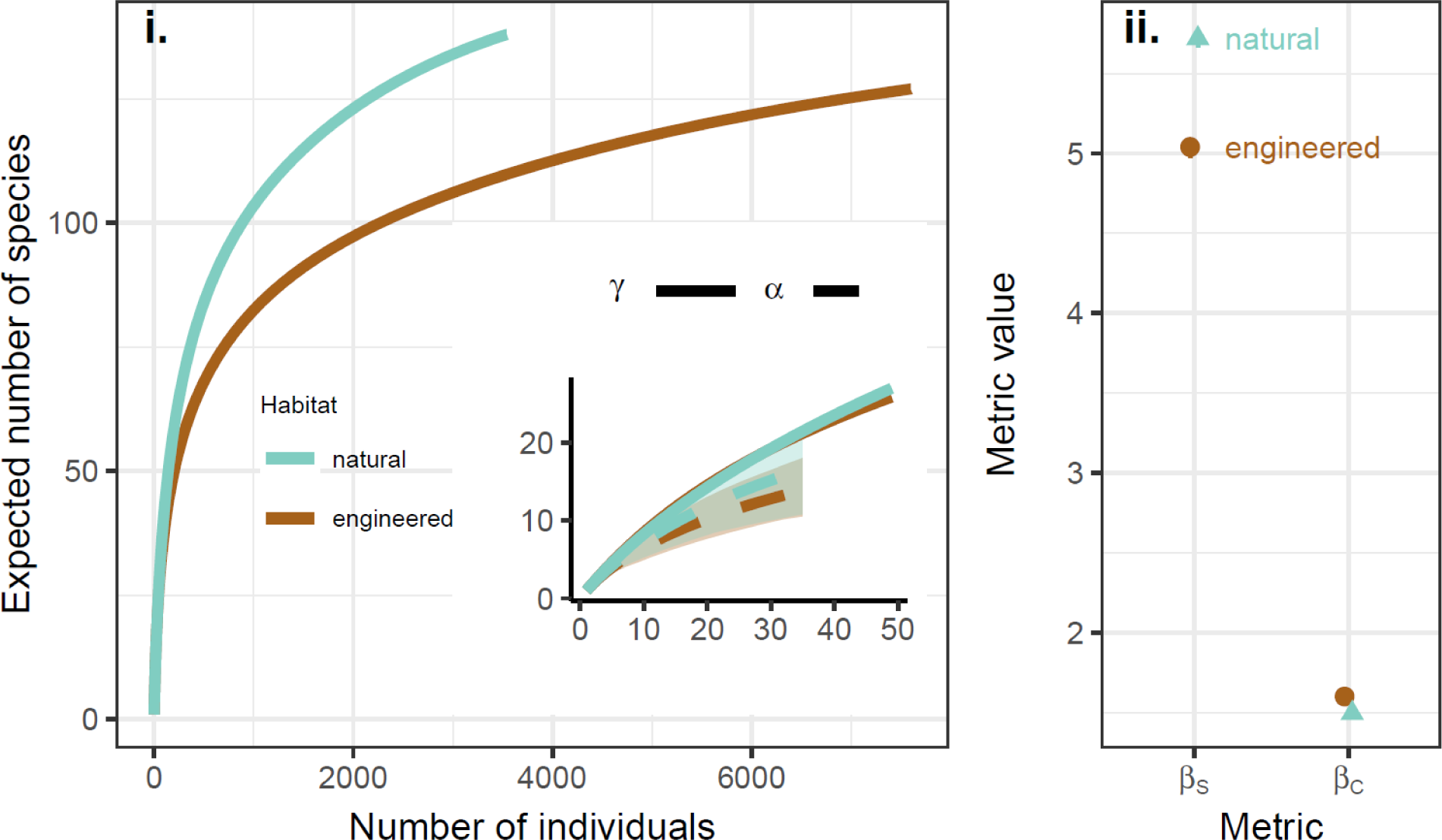
Total spatiotemporal *β*-diversity: (i) *γ*-scale rarefaction curves for each habitat, with inset showing *γ*- and average *α*-scales (note an individual α-scale curve was a single site in a single year); (ii) mean *β_S_* and *β_C_* [point] (95% quantile whiskers not visible) of total spatiotemporal *β*-diversity jackknife resamples in each habitat type.

#### Q2) How does spatial variation in community composition change through time in each of the two habitats?

Figure 5 shows the pattern of spatial β-diversity in engineered and natural habitats through time. β*_S_* increased through time for the natural sites, indicating that those communities were becoming more different from one another through time (opposite to the oft expected pattern of biotic homogenization, where communities become more similar through time and spatial β diversity declines). There was no similar trend in spatial β*_S_* of the engineered sites, and by the end of the time series (but not the beginning) the engineered sites had lower levels of β*_S_* than the natural sites. However, this pattern qualitatively changed when the influence of non-random patterns was explicitly considered. β_C_ indicates that species became less aggregated within engineered sites through time, suggesting biotic homogenization after random-placement mechanisms were controlled, and no change in β_C_ in the natural habitat. Combined, these results suggest the apparent pattern of increasing differentiation in the natural habitat was mostly driven by sampling effects (e.g., altered numbers of individuals, and/or rare species), and that there was a weak decrease of within species aggregation across sites in the engineered habitat.

**Figure 5.**
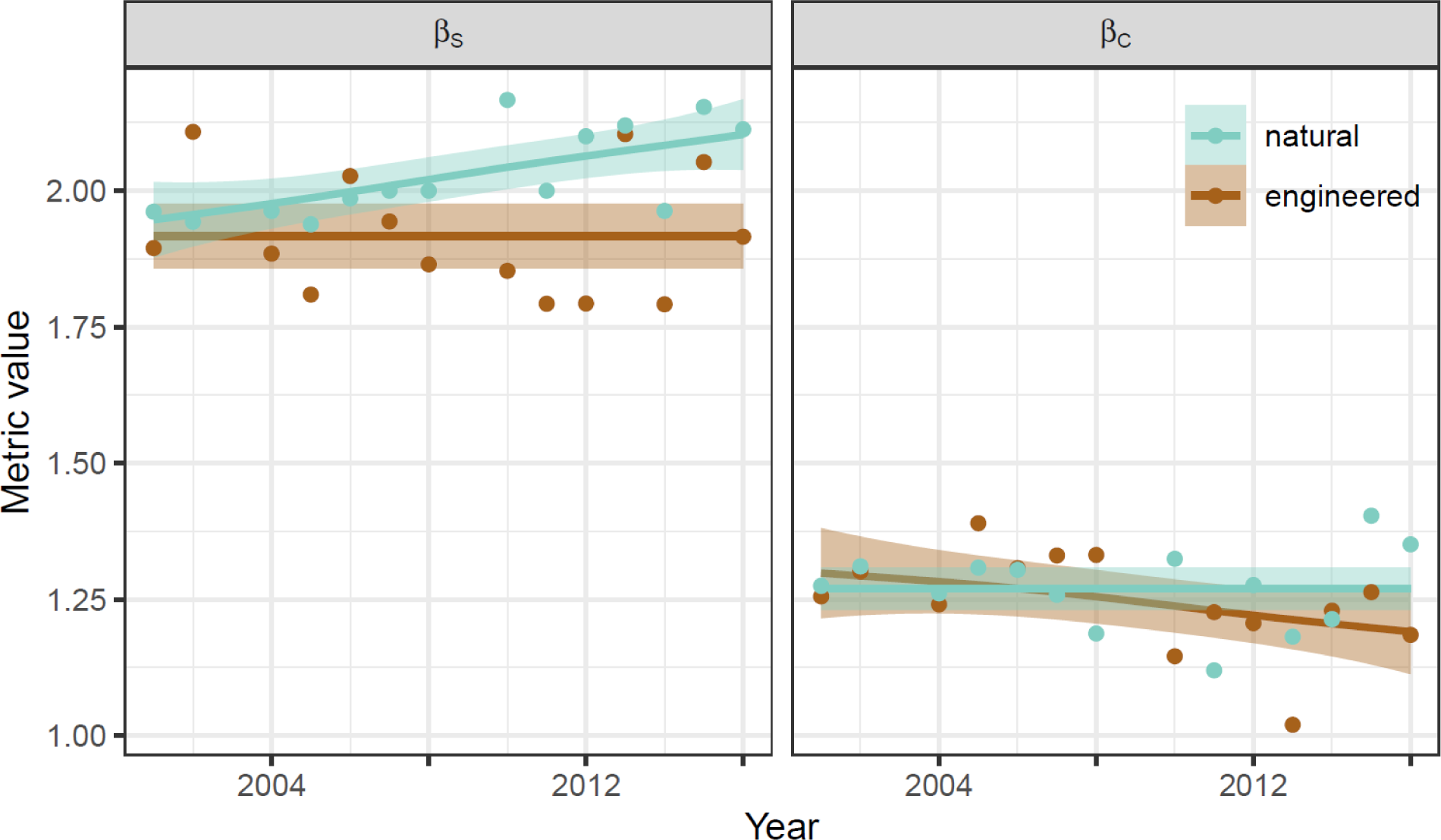
Spatial *β*-diversity as a function of time for *β*_S_ and *β*_C_ in the two habitats. Trend lines represent OLS linear models with their 95% CI.

#### Q3) Does the temporal variation in community composition differ between engineered and natural habitats?

Across all years, sites in the engineered habitat had greater variation than sites in natural habitat in both the total number of individuals (i.e., the end points of the γ-scale curves on the x-axis, Fig. 6i), and shape of the SAD (reflected by greater variation in the curvature of the γ-scale rarefaction). On average, natural sites had slightly higher levels of temporal β*_S_* than the engineered sites, but the variation among sites (and only three replicates) meant there was no overall difference in temporal β*_S_* between habitats (Fig. 6ii). We conclude that the weak differences of temporal β*_S_* between habitats were primarily due to random sampling effects (Fig. 6i) because this pattern disappeared for β*_C_* (and β*_C_* was slightly higher in the engineered habitat). In both habitats, β*_S_* was also more than double the value of β*_C_*, suggesting that more than 50% of year-to-year variation in community composition was due to changes in the number of individuals and/or rare species. The similar values β*_C_* in both habitats indicates that temporal autocorrelation of species presences did not differ much between natural and engineered habitats.

**Figure 6.**
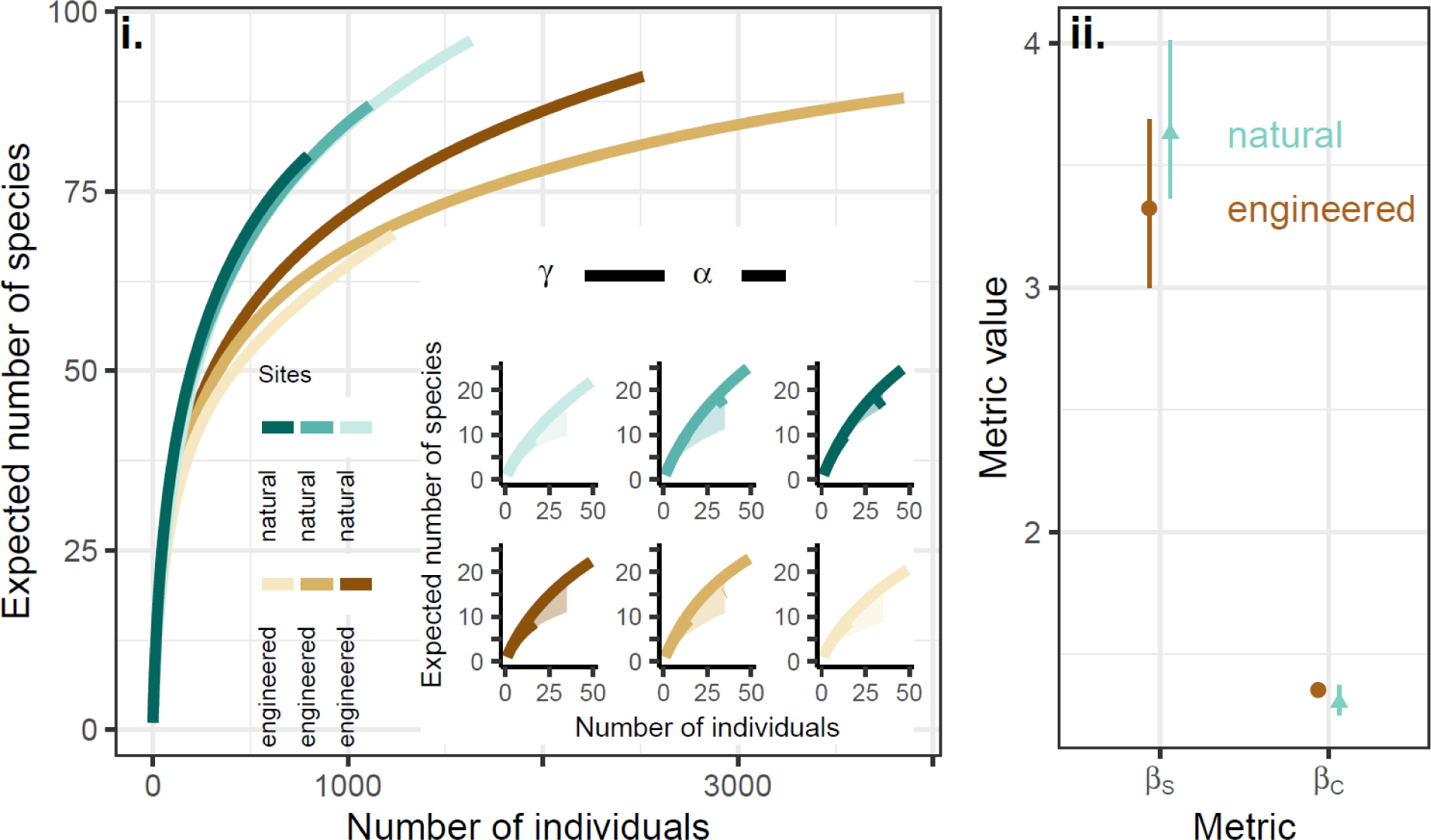
Temporal *β*-diversity: (i) γ-scale rarefaction curves for each site (i.e., all years combined), with inset showing *γ*- and the average α-scales for each site; (ii) temporal *β*_S_ a_nd_ *β*_C_ (mean [point] and 95% quartiles [whiskers] of jackknife resamples) of total temporal *β*-diversity.

#### Q4) Are there compositional differences between engineered and natural habitats, and do any differences change through time?

Finally, the same concepts and tools that we used to examine variation in species composition within treatments can also be used to compare species composition between treatments through time. Essentially, this asks whether bird communities in natural and engineered sites are random subsets of a common larger species pool? Do non-random spatial patterns contribute to any differentiation? And do these patterns change over time? (Fig. 7). Here, the overall difference between treatments (β*_S_*) was larger than one (there is some species turnover between habitats) and slightly declined through time (homogenization). However, when only non-random patterns were considered, β*_C_* was closer to, though still greater than one, and only slightly declined through time. This suggests that once we control for sampling effects that compositional differences between the habitats were relatively small but still detectable and not changing through time.

**Figure 7.**
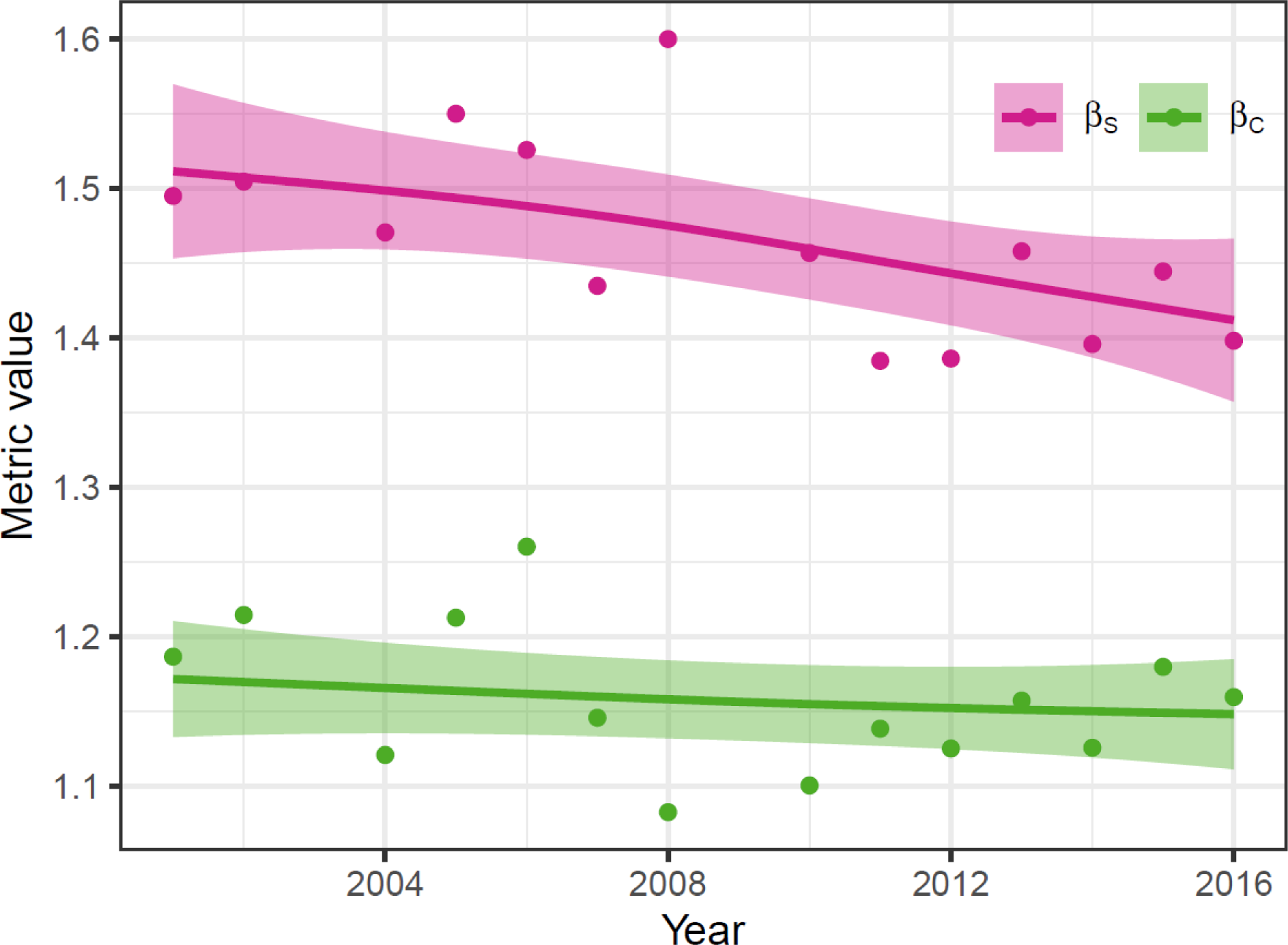
Compare species composition between the engineered and natural riparian habitat types (i.e., the *β*-diversity) as a function of time.

## Discussion

We have described and demonstrated an integrated framework for quantifying the underlying causes of β-diversity and namely if those causes are due to random sampling effects or aggregating mechanisms such as environmental filtering and dispersal limitation. The approach that we have described provides a generalized framework for comparing patterns of total β-diversity, which we call β_S_, to those that specifically partial out the non-random patterns of β-diversity (β_C_). We have demonstrated that any question relating to how species composition changes across samples, whether they be taken through space or time, can be subject to the same approach and metrics. For many cases, this greatly simplifies what can seem a complex endeavor of finding the ‘right’ β-metric for the question at hand.

Often, researchers switch β-diversity metrics and concepts when measuring compositional shifts within a metacommunity or among heterogeneous sites along an environmental gradient (e.g., Anderson et al. 2011). For example, within a metacommunity, estimates of β-diversity are often based on measures of dispersion in community composition among sites (e.g., Anderson et al. 2006). These measures, however, can be strongly influenced by both the relative abundances of species, and the size of the regional species pool. This means that randomization-based null models) are needed if one wants to compare levels of dispersion among different metacommunities, and/or make inferences regarding potential driving mechanisms (e.g., Chase et al. 2011, Kraft et al. 2011, Myers et al. 2013). However, the appropriate form of randomization for the null model remains contentious (Kraft et al. 2012, Qian et al. 2012, Mori et al. 2015, Tucker et al. 2016, Xing and He 2021). Our rarefaction-based approach can also be considered a type of null sampling model. However, comparing rarefaction curves has a number of benefits over other null model approaches: by calculating α- and γ-scale curves, β-diversity can be put back into the context of scale-dependent multicomponent changes in diversity (Chase et al. 2018, Blowes et al. 2022, Rolls et al. 2023); rarefaction curves can be based on analytical solutions improving efficiency; and, rarefaction curves can be visualized, making them more intuitive and easier to communicate than other null model approaches. Nevertheless, some of the concerns arising from the use of null models also apply to the approach overviewed here. For example, there is a strong ‘Narcissus’ effect (i.e., the outcome reflects the inputs) in developing null models to evaluate whether differences among samples deviate from a null expectation; the samples that are used to calculate γ-diversity influence the likelihood that they will deviate from a null expectation (Ulrich et al. 2017). The same is certainly true for the use of individual-based rarefaction curves in which deviations are mathematically constrained by the two end points of the rarefaction curve (McGlinn et al. 2021).

Baselga (2010) has advocated an approach that partitions measures of dissimilarity among samples (e.g., Jaccard’s or Sorensen’s index or an abundance-based equivalent) into measures that capture species turnover between samples, and those that account for the nestedness of species difference between samples (but see Šizling et al. 2022). In essence, the nestedness part of this partition is the same as our ‘random-placement’ effect, while turnover captures the essence of our β-diversity measures that capture non-random variation among samples. For example, in our case study we asked whether bird species in engineered and natural riparian habitats were a random subset of the same regional species pool (Figure 7). We found that β_S_ values were quite high compared to the β_C_ values, which indicates the turnover component is small relative to the nested component in Baselga’s approach.

As with spatial β-diversity comparisons, there have been variable approaches to capture β-diversity through time (Legendre 2019, Magurran et al. 2019, Tatsumi et al. 2022). Most measures of temporal turnover calculate turnover as a metric of community dissimilarity through time. Often rates of change between an initial and subsequent samples, or the rate of decay in dissimilarity as a function of the time elapsed between samples being compared are estimated, which can then be compared across systems or taxa (e.g., Korhonen et al. 2010, Blowes et al. 2019). However, as with spatial β-diversity, these measures cannot discern whether observed rates of turnover are different from what would be expected from a random placement model through time. Authors have used different approaches to remedy this problem. For example, Dornelas et al. (2014) compared rates of temporal β-diversity to those expected from a neutral model (Hubbell 2001) to discern whether turnover rates were faster than expected under the assumption of neutral dynamics, while Stegen et al. (2013) used a null model to determine whether temporal turnover patterns were greater than expected from sampling effects. Temporal turnover can also be decomposed into changes due to abundances (similar to our ‘sampling’ effects) and changes due to species turnover (Shimadzu et al. 2015, Lamy et al. 2015). As with spatial β-diversity measures, our approach is similar, but simplifies the problem by asking whether temporal changes are non-random in a time series.

Recently, authors have developed approaches to partition the influence of species gains and losses to changes in spatial β-diversity through time (Rosenblad and Sax 2017, Tatsumi et al. 2021), and these have been expanded to incorporate changes in relative abundances (Tatsumi et al. 2022). These methods are useful for examining “winning” and “losing” species that underlie changes in spatial β-diversity through time. However, these methods risk isolating beta-diversity changes from local (*α*) and regional (*γ*) scale changes, and are unable to disentangle random versus non-random structure associated with these changes. Thus, our approach can provide a complementary, and more complete picture into scale-dependent changes driving variation of spatial composition through time.

Finally, for simplicity we have focused here on two related metrics: β_S_ and β_C_. Other measures of β diversity with different weights on common and rare species (i.e., Hill numbers) (Jost 2007, Tuomisto 2010, Chao et al. 2012, 2023, but see Lande 1996) can also be calculated at different points along the rarefaction curves. For example, the metric based on Simpson’s entropy, also known as the probability of interspecific interaction (PIE) (or Gini-Simpson index)(Hurlbert 1971) (where *q*=2 in the Hill number continuum; Jost 2007, Chao et al. 2014), can be visualized as the slope at the base of the rarefaction curve (Chase et al. 2018, McGlinn et al. 2019). These Hill numbers or numbers equivalents can also be used with the multiplicative diversity partition used here (i.e., *γ* = *α*\**β*; Jost 2007), and result in an effective number of distinct communities, with the tuning parameter (i.e., order *q*) determining the sensitivity to rare and common species. Recently, Chao et al. (2023) also proposed a framework for standardizing beta-diversity that consider the joint influence of sampling effects and spatial/temporal aggregation. Their framework and ours both standardize biodiversity data to the same level of sample coverage when comparing β between meta-communities. However, an important difference between the approaches is that Chao et al. (2023) assume that individuals are independently sampled (i.e., randomly encountered), whereas we assume that individuals within a sample are not independent of one another due to aggregation. In fact, our primary intention here is to explicitly quantify the important contributions of aggregation to β-diversity, which cannot be directly measured with the Chao et al. (2023) approach.

## Conclusions

Ecologists are often interested in examining the role of metacommunity-level mechanisms such as dispersal limitation and environmental filtering for patterns of β-diversity (Vellend 2016, Leibold and Chase 2018). The generalized approach that we have described relies on a set of intuitive metrics from sampling theory to quantify total β-diversity (*β*_S_), and *β*-diversity due to non-random aggregation (*β*_C_), which will allow for stronger tests of hypotheses related to mechanisms expected to influence patterns of aggregation. In addition, the framework provides an integrated way to examine how changes at finer (*α*) and coarser (*γ*) scales combine to determine variation in species composition (*β*). This places a central focus on scale-dependent diversity changes, with the potential to uncover deeper insights into scale-dependence by varying the focal spatial or temporal grain of the analysis. It remains an open question as to how much variation in β-diversity reflects random sampling effects vs non-random aggregation effects. Our framework provides a means of addressing this question across space and time.

## Acknowledgements

DJM, SAB, TE, JMC gratefully acknowledge the support of the German Centre of Integrative Biodiversity Research (iDiv) Halle-Jena-Leipzig (funded by the German Research Foundation; FZT 118). ISM received funding from the European Union Horizon 2020 research and innovation programme under the Marie Sklodowska-Curie grant agreement no. 894644. TE was supported by the German Research Foundation (DFG) within the project “Establishment of the National Research Data Infrastructure (NFDI)” in the consortium NFDI4Biodiversity (project number 442032008).

